# Extreme genome scrambling in cryptic *Oikopleura dioica* species

**DOI:** 10.1101/2023.05.09.539028

**Authors:** Charles Plessy, Michael J. Mansfield, Aleksandra Bliznina, Aki Masunaga, Charlotte West, Yongkai Tan, Andrew W. Liu, Jan Grašič, María Sara del Río Pisula, Gaspar Sánchez-Serna, Marc Fabrega-Torrus, Alfonso Ferrández-Roldán, Vittoria Roncalli, Pavla Navratilova, Eric M. Thompson, Takeshi Onuma, Hiroki Nishida, Cristian Cañestro, Nicholas M. Luscombe

## Abstract

Genes are not randomly distributed throughout chromosomes. How gene order evolves and how selective constraints act to preserve or vary gene order, both at the macrosyntenic level of whole chromosomes or microsyntenic level of gene blocks, are central questions of evolutionary biology and genomics that remain largely unsolved. Here, after sequencing several genomes of the appendicularian tunicate *Oikopleura dioica* from different locations around the globe, we show an unprecedented amount of genome scrambling in animals with no obvious morphological differences, consistent with cryptic speciation. Our assemblies suggest that all members of this clade possess a common 3-chromosome karyotype, and that different species largely preserve gene content, despite the presence of thousands of rearrangements in gene order. The movements of genes are largely restricted to chromosome arms and sex-specific regions, which appear to be the primary unit of macrosynteny conservation, and examples of these within-arm movements can be seen in the *Hox* and *Fgf* gene families. Our approach employing whole-genome alignments demonstrates that segments containing protein-coding elements tend to be preserved at the microsyntenic scale, consistent with strong purifying selection, with appreciably less preservation of non-coding elements. Unexpectedly, scrambling did not preserve operon structure across species, suggesting an absence of selective pressure to maintain operon structure. As well, genome scrambling does not occur uniformly across all chromosomes, as short chromosome arms possess shorter genes, smaller operons, more breakpoints, and elevated dN/dS values compared to long chromosome arms. Estimation of divergence times among the cryptic *O. dioica* lineages yielded an estimated breakpoint accumulation rate of 6 to 25 breakpoints per megabase per million years, which is an order of magnitude higher than the rates for other ascidian tunicates or *Drosophila* species. Therefore, *O. dioica* appears to be an attractive animal system to unravel the mechanisms that underlie gene order and synteny conservation, as well as exploring the limits of genome scrambling without an apparent impact on phenotypic evolution.

## INTRODUCTION

It is widely accepted that the distribution and order of genes on chromosomes is not random, as changes in gene order are likely to affect the regulation of gene expression. However, how evolution acts to preserve or vary gene order remains poorly understood. Comparisons of distantly-related groups of metazoans have revealed gene linkages within chromosomes that have been preserved across many lineages for more than half a billion years (Simakov et al. 2022). The conservation of gene linkage is a feature referred to as “conserved **synteny**”, from the Greek meaning “same ribbon”, which describes homologous genes that co-locate within a single chromosome (Passarge, Horsthemke, and Farber 1999). Differences in the scale and extent of synteny conservation have led to the concepts of “**micro**-” and “**macro**-” synteny. Microsynteny (also known as “**collinearity**” in genomics) refers to the conservation of gene content and order within sets of tightly-linked orthologous genes. Generally, closely-related species tend to possess greater conservation of microsynteny, and for this reason, it can be used to clarify phylogenies (Drillon et al. 2020; Pereira-Santana et al. 2020). While microsynteny is generally weakly conserved in distantly-related species, the remnants of ancient linkage karyotype groups can be detected at chromosome scale; the conservation of genes on chromosomes that can be traced back to an ancestral karyotype is reflected in the concept of macrosynteny, examples of which include the chromosomal conservation that can be traced back the last common ancestor of metazoans (Simakov et al. 2022). Strikingly, there are many notable examples of conserved microsynteny even among distantly-related organisms. The most famous example of conserved microsynteny in animals is the *Hox* cluster, which contains genes that regulate axial patterning during embryogenesis, and whose ancestry can be traced back to the origin of bilaterian animals hundreds of millions of years ago (see Wanninger 2023 for a recent review). There are many other examples of highly conserved microsyntenies across metazoans, in many cases related to the functional constraints imposed by cis-regulatory elements on the coordinated transcription of nearby genes. This includes genomic regulatory blocks (GRBs), within which the action of conserved non-coding elements allows the coordinated expression of genes in a local genomic neighbourhood (Hurst, Pál, and Lercher 2004; Rowley and Corces 2018; Engström et al. 2007; Irimia et al. 2012). Thus, evaluating the conservation and loss of synteny can provide important information for generating testable hypotheses related to gene regulation, genome biology, and evolution.

The loss of synteny can be provoked by genome rearrangements, such as chromosome translocations related to unequal recombination, or chromosome fragment mobilisation due to transposon activity. Both of these processes can result in changes of gene order and the reallocation of genes to different neighbourhoods. The accumulation over time of many rearrangements over time results in genome “scrambling”, a concept that in linguistics refers to language syntaxes that permit changes in word order without altering the meaning of a sentence. Scrambling has been used to describe the patterns of synteny loss in genomic comparisons of distantly-related species, such as fugu and humans, whose genome organisation has significantly diverged over hundreds of millions of years (Aparicio et al. 2002). However, fundamental questions remain, such as how evolutionary forces act to constrain or accelerate the rate of rearrangement, or how phenotypic differences could be related to rearrangements. Addressing these problems is difficult at large time scales and genetic distances.

In order to better understand the phenomenon of genome scrambling and how it is shaped by evolutionary forces, here we study the zooplanktonic appendicularian tunicate *Oikopleura dioica.* Our recent work revealed that different populations of *O. dioica*, despite lacking any obvious morphological distinction, indeed represented multiple cryptic species. Populations sampled from the Japanese Seto inland sea (Osaka University laboratory strain) and from the sub-tropical island of Okinawa, Japan (OIST laboratory strain), were shown to be reproductively isolated and exhibited remarkably high genetic distance despite their overall phenotypic similarity (Masunaga et al. 2022). The process of generating a telomere-to-telomere assembly and gene annotation of *O. dioica* from Okinawa (Bliznina et al. 2021) further prompted important differences in gene organisation through comparison with the assemblies of individuals from Osaka (Denoeud et al. 2010; K. Wang et al. 2020; Bliznina et al. 2021) and Bergen (Norway), suggesting that *O. dioica* species around the globe could provide an attractive system to study the loss of conserved synteny in the absence of phenotypic differences. *O. dioica*’s karyotype comprises 3 chromosome pairs (Körner 1952; Liu et al. 2020): two acrocentric autosomes and an acrocentric sex chromosome containing a pseudoautosomal region common to the X and Y chromosomes (Denoeud et al. 2010). The Y-specific region is repeat-rich, gene-poor, and differs from all other genomic regions. The pseudoautosomal region contains the sex chromosome’s centromere and long-read sequencing suggests that it is connected to sex-specific regions by a ribosomal DNA locus (Bliznina et al. 2021). Chromosome contact analysis suggests that there is relatively little interaction between the arms of individual chromosomes or sex-specific regions, similar to the “type-I” genome architecture reported by Hoencamp et al. (2021).

*O. dioica* has the smallest reported non-parasitic animal genomes reported to date (Denoeud et al. 2010; K. Wang et al. 2020; Bliznina et al. 2021; H. C. Seo et al. 2001). This genome reduction appears to be the result of a drastic process of compaction involving a reduction in repeat content (∼15%) (Henriet et al. 2015), numerous gene losses (see (Ferrández-Roldán et al. 2019) for a recent review), and the evolution of numerous operons (Danks et al. 2015; Ganot et al. 2004). In *O. dioica*, a significant fraction of genes are densely packed in a microsyntenic head-to-tail configuration and transcribed in polycistronic mRNAs, which are processed by the addition of a *trans*-spliced leader RNA (Ganot et al. 2004). In contrast to bacterial operons, where co-transcribed genes tend to be functionally related, in *Oikopleura* and other animals, the functions of genes in operons are more loosely related, with a trend towards house-keeping, cell cycle, translation and germline functions. How these co-transcriptional units, which likely evolved independently in different eukaryotic lineages, is unknown (Zeller 2010; Danks et al. 2015; K. Wang et al. 2015). At the same time, genome compaction in *O. dioica* also appears to have been accompanied by a drastic loss of conserved microsynteny compared to other chordate genomes, including the disintegration of the paradigmatic *Hox* cluster (H.-C. Seo et al. 2004).

Here, we report and examine the phenomenon of genome scrambling in *O. dioica* by comparing chromosome-scale genome assemblies from *O. dioica* from different locations in the Atlantic and Pacific oceans in the Northern Hemisphere. In spite of their broadly conserved morphology and karyotype, these populations exhibit a considerable degree of genetic rearrangement. These rearrangements tend to preserve coding elements within the genome, with exons exhibiting stronger conservation than genes, and operons exhibiting little conservation overall. Further, there is heterogeneity of conservation within individual chromosomes, such that short chromosome arms accrue proportionally greater point mutations and breakpoint regions. Remarkably, these genetic events appear to have accrued much more quickly in *O. dioica* than other animals.

## RESULTS

### A pan-genomic analysis of *Oikopleura dioica*

We have generated two high quality chromosome-scale genome assemblies of *O. dioica* from individuals from the Mediterranean Catalan coast (“Barcelona”, Supplementary Table 1) and from the Osaka University laboratory strain (“Osaka”, Supplementary Table 1), which are added to our previously reported telomere-to-telomere genome assembly from an individual from the coastline of the OIST laboratory strain (“Okinawa”, Supplementary Table 1) (Bliznina et al. 2021). We validated each reference assembly using an additional contig-level assembly from the same species (respectively “Bergen“, “Aomori“, “Kume“, Supplementary Table 1), yielding one chromosome-scale genome sequence and one contig-level assembly per species (Supplemental data 1). We assembled the Barcelona genome using a similar procedure to the Okinawan genome, including the use of chromosome conformation information (Hi-C libraries) to aid scaffolding. The Hi-C contact map showed that the chromosome arms and the sex-specific regions had few interactions with each other, and the assembly graph connected the sex-specific regions to the pseudoautosomal region’s long arm through ribosomal DNA repeats. Moreover, we have constructed a new Osaka genome assembly by scaffolding the OSKA2016 assembly (K. Wang et al. 2020) with long Nanopore reads that we sequenced from single individuals from the same laboratory strain. To ensure consistency (Weisman, Murray, and Eddy 2022), we generated updated annotations for all genomes using a common automated pipeline, including repeat masking and gene prediction steps, which provide a highly robust set of annotations that facilitate inter-species comparisons (Supplementary Table 1). Analysis of the chromosomal configuration of Barcelona, Osaka and Okinawa revealed a common conserved karyotype consisting of 3 chromosome pairs: two acrocentric autosomes and an acrocentric sex chromosome containing a sex-specific region (XSR or YSR) and a pseudoautosomal region (PAR) common to the X and Y chromosomes.

### The scrambled genomes of *Oikopleura dioica*

To investigate the evolution of the chromosomes in *O. dioica* from Okinawa, Osaka and Barcelona, we first performed pairwise whole genome alignment with LAST, which is especially good at finding structural rearrangements and recombinations (Frith and Kawaguchi 2015; Mitsuhashi et al. 2020). Coupling the LAST method and the Nextflow pipeline system (Di Tommaso et al. 2017), we developed a reproducible, standardised method to identify sequences with one-to-one correspondence (mapping) between pairs of genomes. The computation of all-by-all pairwise genome alignments revealed an unprecedented level of genetic rearrangement for closely-related cryptic species (Figure 1A). For example, the line plot comparing the whole genome sequences of *O. dioica* from Osaka and Okinawa revealed a striking pattern, with little to no conservation of collinear DNA segments on any chromosome, consistent with an extreme degree of scrambling across the entire genome (Figure 1A).

**Figure 1:**
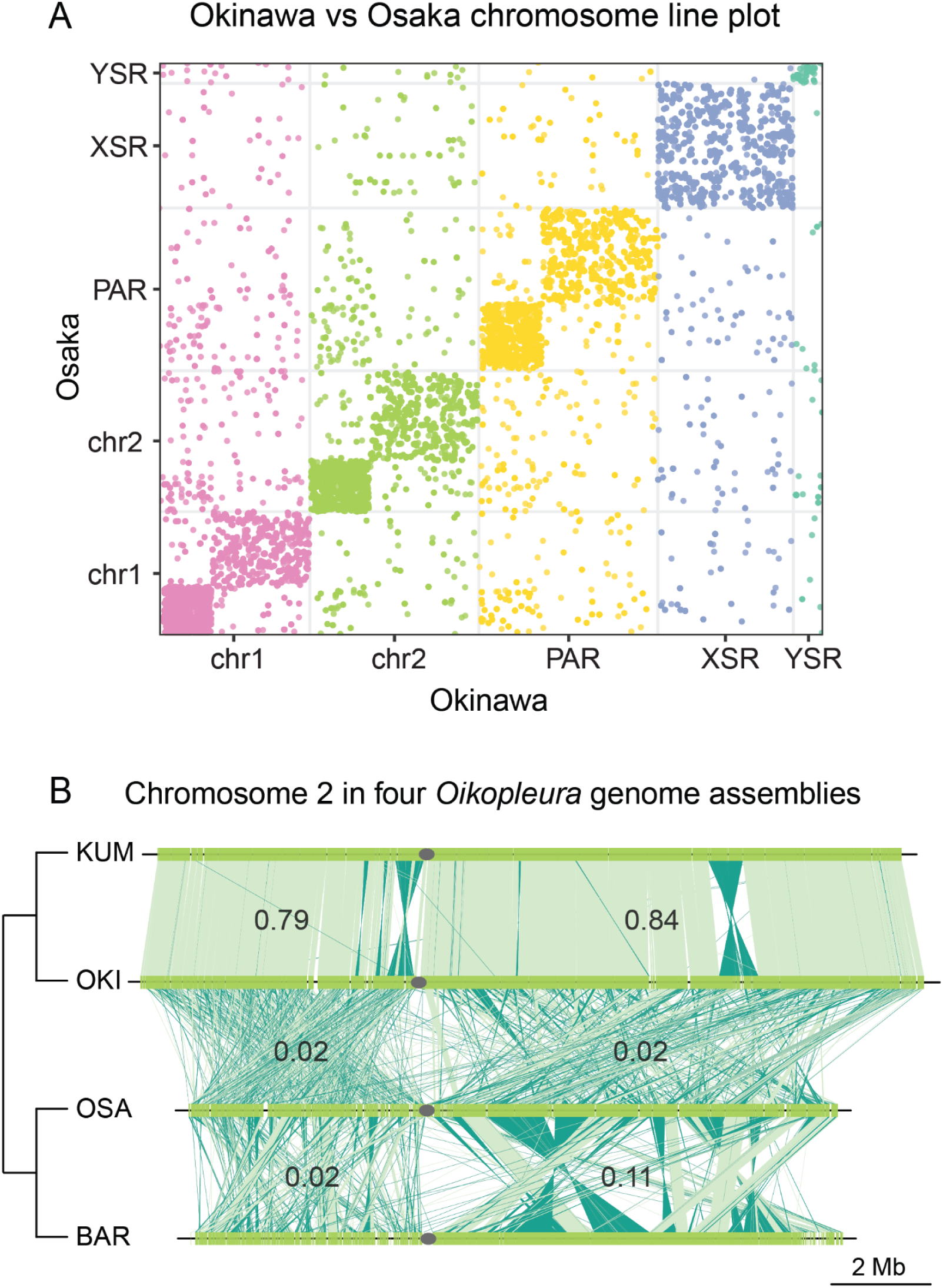
Extensive genomic rearrangement has occurred between cryptic *Oikopleura dioica* species. A) Line plot representation of the pairwise whole-genome alignment between Okinawa and Osaka genomes. Significant genomic rearrangement occurs across the length of all chromosomes. B) Pairwise comparisons of chromosome 2 between *Oikopleura* genomes. Top to bottom: Kume, Okinawa, Osaka, Barcelona. Dark green indicates plus/minus strand alignments and the grey ellipse the centromere. The cladogram is deduced from the scrambling index (Table 1).

Multi-chromosome line plots comparing *O. dioica* from different locations further revealed that the genome scrambling phenomenon was not restricted to the comparison of Osaka and Okinawa, but instead common amongst all compared genomes (Figure 1B). In general, the extent of genome scrambling increased with increasing genetic distance, as measured by previous analyses of marker genes (Masunaga et al. 2022). Comparisons of *O. dioica* originating from the same geographic region, such as the comparison of two individuals from the subtropical North Pacific population in Okinawa and Kume, showed little scrambling, with large, intact collinear segments of DNA visible. Comparable levels of scrambling were observed for within-region comparisons of Osaka and Aomori from the North Pacific, and Barcelona and Bergen from the North Atlantic. However, much more scrambling was evident in comparisons between geographic regions, such as the comparison between Osaka and Barcelona (Figure 1B). These observations suggested that genome scrambling was therefore a common evolutionary characteristic in *O. dioica* genomes.

To quantify the degree of scrambling in a pair of genomes and determine how scrambling might relate to other measures of genetic distance, we created a “scrambling index”, which measures the degree of strand randomisation, and thus the loss of collinearity, between aligned regions. A scrambling index value between 0.5 and 1 indicates that most alignment strands align in the same orientation (i.e., plus-to-plus or minus-to-minus); scrambling index values approaching 0 indicate that either alignment orientation is equally frequent (i.e., plus-to-minus and vice-versa). Computation of the scrambling index for each genome pair (Table 1) yielded high values for within-region comparisons (Okinawa vs. Kume, Osaka vs. Aomori, and Barcelona vs. Bergen), consistent with a lesser degree of scrambling and rearrangement in these comparisons. Very small scrambling indices were obtained for comparisons of the subtropical North Pacific to other groups, indicating an extreme degree of scrambling. Interestingly, comparisons between animals from the temperate North-Pacific and North-Atlantic locations also yielded low SI values (near 0.2), which is congruent with the relatively high level of scrambling observed in line plot comparisons of Osaka and Barcelona (Figure 1B). Considering that the large amount of genome scrambling between Osaka and Barcelona is likely to compromise chromosome pairing during meiosis, we interpreted it as an indication that *O. dioica* from those distant locations are likely to be reproductively isolated and therefore considered different cryptic species. In summary, our results suggest that the study of genome scrambling in *O. dioica* around the globe can be useful to discover additional cryptic species and assess its relationship to reproductive isolation, thereby illuminating the evolutionary history of this cosmopolitan species.

**Table 1:** Scrambling index for same-species (green), North Atlantic–North Pacific (yellow), and subtropical North Pacific–other (red) pairs of genomes. A value of 1 indicates complete strand-identity, while 0 indicates complete strand randomization. Importantly, our contig assembly set (Kume, Aomori and Bergen) allowed us to rule out technological biases introduced by different sequencing technologies as a potential cause underlying the identified loss of synteny, as high within-species scores were obtained with three independent combinations of sequencing platforms. The contig-level assemblies also provided further validation through independent comparisons of species pairs (for instance, the scrambling index value for the Osaka-Bergen and Aomori-Barcelona comparisons are similar). However, since contig-level assemblies (Kum, Aom, and Nor) are not scaffolded to the chromosome-scale or oriented from telomere-to-telomere, their scrambling index values are penalised, trending towards 0.5 for comparisons with their chromosome-scale counterparts (Oki, Osa, and Bar, respectively).

|  | Oki | Kum | Osa | Aom | Bar | Nor |
| --- | --- | --- | --- | --- | --- | --- |
| Oki | 1 | 0.675 | 0.045 | 0.08 | 0.1 | 0.18 |
| Osa | 0.045 | 0.06 | 1 | 0.545 | 0.23 | 0.285 |
| Bar | 0.1 | 0.115 | 0.23 | 0.22 | 1 | 0.5 |

### Impact of genome scrambling on macrosynteny conservation in *O. dioica*

Line plots between Osaka and Okinawa showed that the rearrangements causing genome scrambling were largely restricted to homologous chromosomes (∼94% of all rearrangements were intrachromosomal), while interchromosomal rearrangements were rare (Figure 1A). Within each chromosome, rearrangements tended to occur within arms or the sex-specific regions (∼99%). Thus, the primary unit of macrosynteny conservation among *O. dioica* species appears to be chromosome arms and sex-specific regions (which for the sake of simplicity we will also refer to as “arms”). To investigate the impact of genome scrambling on the evolution of synteny blocks, we compared gene order conservation across species. We computed 5162 groups of single-copy orthologues present in the 6 genomes, and visualised them with strand-independent macro-synteny dotplots, which show the positions of the same gene in a pair of genomes (Figure 2A). Similar to whole-genome nucleotide alignments (Figure 1), the gene-based dot plots confirmed that gene order rearrangements were mostly restricted to homologous arms (Figure 2A). As well, this analysis of orthology revealed that some interchromosomal translocations observed at the nucleotide level also involved whole-gene translocations between chromosomes. Macrosynteny dot plots also revealed that large synteny blocks may be conserved in comparisons of the same species (e.g. temperate North-Pacific Osaka vs Aomori), with less conservation evident in comparisons between distant locations (e.g. Osaka vs Barcelona), and nearly undetectable in comparisons of Okinawa versus Osaka or Barcelona. The number of orthologues per synteny block also decreased with increasing genetic distance, with a maximum of 44 for Osaka versus Okinawa, a maximum of 174 for Osaka versus Barcelona, and a maximum of 714 for Osaka versus Aomori (Supplementary Figure S1). Therefore, our data confirmed that genome scrambling in *O. dioica* has resulted in highly variable gene order in these species, and that synteny block conservation decreases with increasing evolutionary distance.

**Figure 2:**
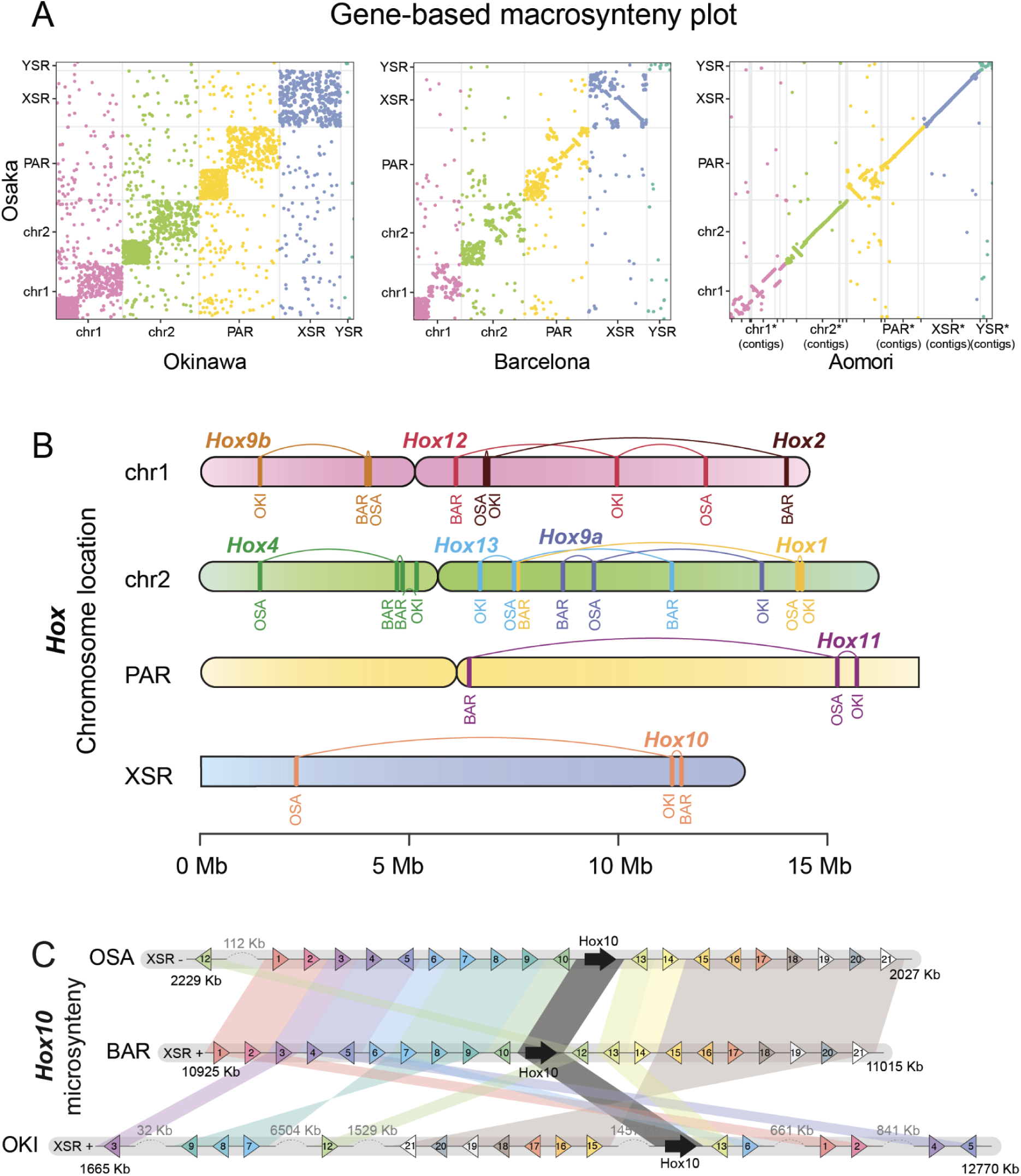
The preservation of orthologous synteny blocks gradually decreases with increasing evolutionary distance in *O. dioica*. A) Dot plots indicating the coordinates of genes belonging to the same orthogroup in pairs of genomes. We used Osaka as a reference genome, as it provided snapshots of genome scrambling at three different stages of the evolutionary process. B) Comparative chromosome mapping of the *Hox* genes in the genomes of *O. dioica* from Osaka (OSA), Barcelona (BAR) and Okinawa (OKI). C) Comparative microsynteny conservation of the block of the next ten genes at each side of the *Hox10* genes in the genomes of *O. dioica* from from Osaka (OSA), Barcelona (BAR) and Okinawa (OKI)

We next mapped the chromosomal locations of several genes associated with characteristic gene clusters (*Hox*, *Fgf*, and *Myosins*) to assess their conservation at the levels of micro- and macro-synteny (Figure 2B, Supplementary Figure S2 and S3). In the case of the *Hox* cluster, while microsynteny conservation has been shown to be essential for embryonic development and axial patterning in vertebrates, the disintegration of the *Hox* cluster in *O. dioica* from Bergen suggested that it may not be essential in this species (H.-C. Seo et al. 2004). Here, we have mapped all *Hox* genes of *O. dioica* in our newly assembled genomes, which corroborated that the full catalogue of *Hox* genes in *O. dioica* is reduced to six genes from which all central *Hox* genes (*Hox3* to *Hox*7) have been lost. Furthermore, we confirmed that the atomization of the *Hox* cluster originally reported in a contig-level assembly in *O. dioica* from Bergen was indeed visible in a chromosome-scale assembly of the analysed species. The presence of *Hox* genes scattered across all chromosomes revealed no trace of the conserved ancestral anterior or posterior macrosynteny of *Hox* genes along chromosome arms (i.e., the anterior *Hox1* and *Hox2* genes are encoded within Chr2 and Chr1; the posterior *Hox10*, *Hox11*, *Hox12,* and *Hox13* genes are encoded on the XSR, PAR, Chr1, and Chr2, respectively; Figure 2B). Comparison of the position of *Hox* orthologues revealed that multiple changes in gene order must have occurred between species, although all have been maintained within the same chromosome arm. Mapping of all six Fgf genes previously reported in *O. dioica* (Oulion, Bertrand, and Escriva 2012) revealed a similar pattern of gene movement, as all gene family members maintain their encoding within chromosome arms (Supplementary Figure S2A). On the other hand, chromosome mapping of the eight Myosin Heavy Chain class II genes presented a different pattern, whereby orthologues occasionally seemed to move more freely including inter-arm and inter-chromosomal translocations (Supplementary Figure S2B). We also inspected patterns of microsynteny in these gene families by examining their 10 nearest neighbouring genes up- and downstream (Supplementary Figure S3). In general, gene families in Barcelona and Osaka exhibited far greater conservation of microsynteny with one another than either does compared to Okinawa. These examples revealed different degrees of microsynteny conservation, ranging from near-complete conservation of entire blocks (e.g. *MyhF*, *MyhG*, *Fgf11/12/13/14a*, *Fgf11/12/13/14b* and *Hox1*) to situations in which a block has seemingly fragmented into many small pieces (e.g. Fgf9/16/20a and *Hox10* or *Hox12*). Thus, the evolutionary pressure to preserve microsynteny likely differs between families, perhaps owing to the nature of the regulatory elements driving their expression. Based on our examination of chromosome mapping for different conserved gene families in *O. dioica*, the position of a gene in one species had surprisingly little predictive power for the position or orientation of that gene in other species.

### Genome scrambling moves short functional regions

A genome alignment in a pair of species can be interrupted because sequence similarity may have eroded over time, or because synteny was truly broken. To investigate the molecular breakpoints responsible for scrambling synteny blocks, we first identified collinear alignments in pairs of genome alignments, defining them as adjacent alignments in the same orientation in both genomes. We termed the regions flanking these collinear alignments “bridge regions”. We then defined “collinear regions” as successions of collinear alignments and bridge regions. By definition, the remaining unmapped regions, for which there is no one-to-one correspondence in a pair of genomes, always correspond to an interruption of collinearity, and we will refer to them as “breakpoint regions”. Lastly, we termed aligned regions that were not collinear to anything as “isolated alignments’’ (Figure 3A). In the comparison of Osaka and Okinawa, isolated regions covered a small fraction of the genome (∼4.7%), while collinear regions (alignments + bridges) covered more than half of it (∼71.8%). Although breakpoint regions tended to be short (0.32 ± 5.1 kbp, *n*=8821, Figure 3D), they covered a considerable fraction of the genome (∼23.5%, Fig 3B), suggesting that the genetic events that produce scrambling are sufficiently ancient for alignability to be lost, or that the mechanism involved the loss of DNA.

**Figure 3.**
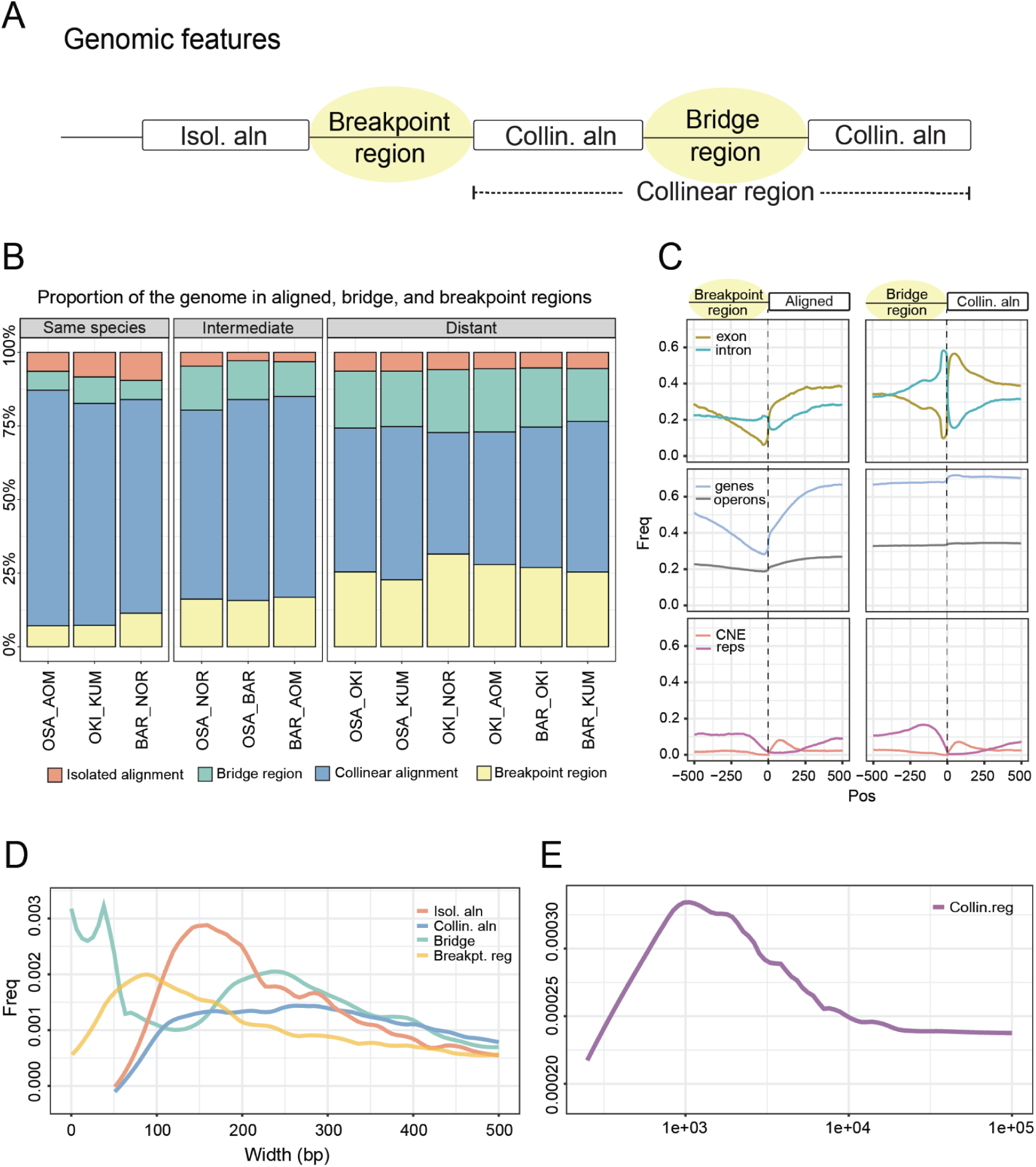
Properties of genomic alignments. A) We divided the aligned and unaligned regions of the genome into four categories according to their participation in collinear regions. Collinear regions are defined as an uninterrupted succession of alignments that are on the same chromosome strand and in the same order in both genomes. B) Proportion of the four categories in different alignment pairs, grouped by evolutionary distance. C) Enrichment of genomic features at the boundary between breakpoint or bridge regions and aligned regions. D, E) Size distribution of the four genome segment categories and of the collinear regions for the Okinawa-Osaka comparison.

To determine how the phenomenon of scrambling related to functional genomic regions, we studied the frequencies of coding and non-coding elements at the boundaries of the four non-overlapping classes of genome segments (Figure 3C). Isolated alignments’ boundaries tended to coincide with exon start positions, and intron stop positions to a lesser extent (Fig. 3C). Isolated alignments were also less frequently part of operons. In terms of non-coding elements, repeat regions were depleted in isolated alignments, while conserved non-coding elements (See Methods) were enriched, with a peak downstream of the alignment start position, consistent with previously reported patterns of erosion (Royo et al. 2011). Breakpoint regions were the least likely to be found within genes. Bridge regions occurred mainly in genic regions, with strong enrichment for introns and repetitive elements (which may be intronic) upstream of collinear alignments; bridge regions were also most frequently associated with operons. The size distribution of segments from each class showed three peaks (Fig 3D): at zero (i.e., a perfectly defined insertion/deletion), at 35 or 36 nt (i.e., average intron size in *O. dioica*), and between 200 and 300 nt. Altogether, the most marked changes in the frequency of genomic elements between classes were related to the frequency of protein-coding features, with the surprising exception of operons, which exhibited modest changes in frequency at the edges of aligned and breakpoint or bridge regions.

### Genome scrambling does not preserve operon structure

Considering the abundance of operons as one of the hallmarks of the *O. dioica* genome (Denoeud et al. 2010), we examined whether operons could impose some limitations to rearrangements in synteny blocks; for example, a single-gene inversion in the middle of a 3-gene operon could result in expression defects by decoupling that gene from its primary regulatory elements. Here, we defined an operon as a set of collinear genes with fewer than 500 base pairs of intergenic space, as this definition produces distributions of operon lengths comparable to the one reported by Denoeud et al. 2010. The number of operons per species ranged between 2379 to 3124 operons, representing between 6653 and 9543 operonic genes, depending on the species (Figure 4A). Unexpectedly, only a small number of operons preserved the same genes across Okinawa, Osaka, and Barcelona (Figure 4A). In line with the larger synteny blocks observed in the comparison of Osaka and Barcelona, a larger number of operons are conserved in this pair (526 operons in common with 350 specific to the pair). Neither Osaka nor Barcelona share a large number of conserved operons with Okinawa, with as little as 176 operons common to all species, suggesting a lack of strong purifying selection to maintain operon structure in *O. dioica*. In agreement with this idea, we found that among protein-coding genetic elements – operons, genes, and exons – operons were the most likely to overlap breakpoint regions. In the Okinawa-Osaka genome pair, 616 out of 1281 operons overlapped a breakpoint (48%), while 5,294 out of 17,291 genes (30%) and 16,787 out of 106,811 exons (15%) overlapped one (Figure 4B). Detailed comparison of operon microsynteny revealed examples of operons with complete conservation located on the same chromosome for Okinawa, Osaka, and Barcelona (Figure 4F). Other examples demonstrate the conservation of an operon following an apparent translocation of some operonic genes to a new location (Figure 4D-E). In some cases, an operon rearrangement involved duplication and translocation of a large portion of an operon into a new chromosome (Figure 4G). While operons are rarely conserved between species in general, operonic genes from one species were significantly more likely to be operonic in a second species across all species-pairs (*p* ≪ 0.001, χ-squared ≫ 4420.2, d.f. = 3). Overall, our data revealed an absence of strong selective constraints to strictly maintain operon structure among species, suggesting operons are prone to be impacted by genome scrambling.

**Figure 4.**
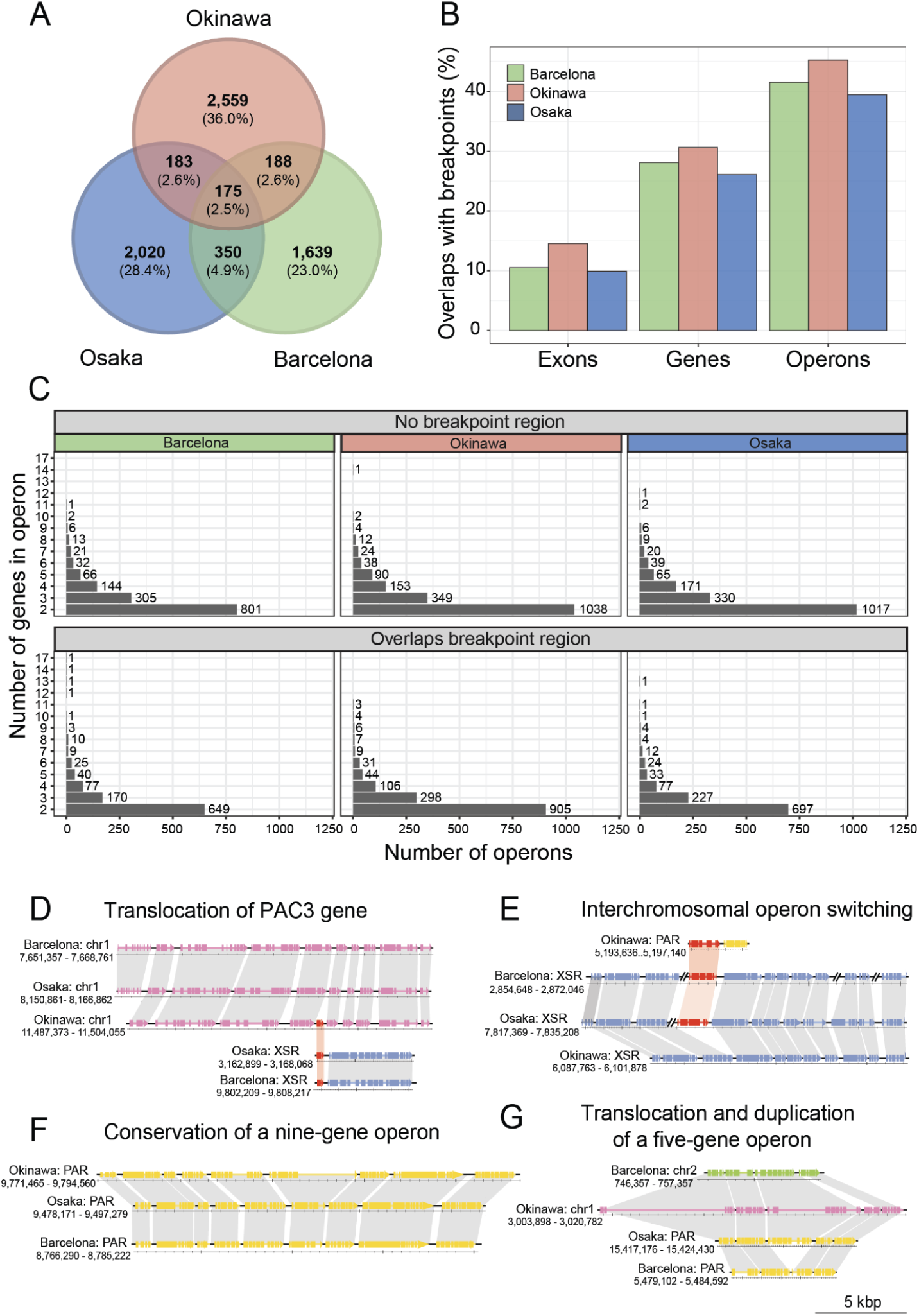
Conservation of operons in the Okinawa and Osaka genomes. A) Most operons are species-specific, as determined by homologous genes. B) The proportion of features that overlap a breakpoint region for each genome. C) Size distribution of operons that overlap or do not overlap a breakpoint region. D) The *PAC3* gene appears to have undergone a translocation between Okinawa and Osaka, changing from one operon to another. E) An example of an operon which is conserved on the XSR of Barcelona and Osaka, in which a central gene appears to have undergone a translocation in Okinawa to a different operon encoded elsewhere in the pseudoautosomal region. F) The 9-gene reported by Ganot et al. is conserved in Osaka, Barcelona, and Okinawa. G) An example of an operon that has been translocated to different chromosomes in each species and duplicated in the Barcelona genome.

### Genome scrambling and chromosomal evolution

Given that the primary scale of macrosynteny conservation among *O. dioica* species appears to be chromosome arms and sex-specific regions, to better understand chromosomal evolution in *O. dioica* species, we investigated the distribution of breakpoint regions, operon sizes and mutation rate at the chromosome level (Figure 5). Our analysis revealed that short chromosome arms consistently exhibited four different qualities than long chromosome arms: first, short arms exhibited a higher relative frequency of breakpoints than long arms; second, short arms contained shorter genes and shorter operons than long arms; third, genes on short arms overlapped breakpoint regions at a higher rate than genes on long arms (∼50% vs ∼20%; *p* ≪ 0.001, χ-squared ≫ 109.6, d.f. = 2); and fourth, dN/dS values were significantly higher on short arms than long arms. In all cases the XSR exhibited patterns comparable to long arms. Our analysis also revealed that these features also consistently varied across chromosome arms, differing between the centres of chromosome arms and subtelomeric or pericentromeric regions. As reported for the Okinawa genome, repeat density increased while gene and operon density decreased in subtelomeric and pericentromeric regions (Bliznina et al. 2021) and Figure Supp 4) for the Osaka and Barcelona genomes. In total, these patterns suggest that in *O. dioica*, short chromosome arms accumulate both point mutations and genome scrambling at a faster rate than long chromosome arms, which leads to longer genes and operons in long arms.

**Figure 5.**
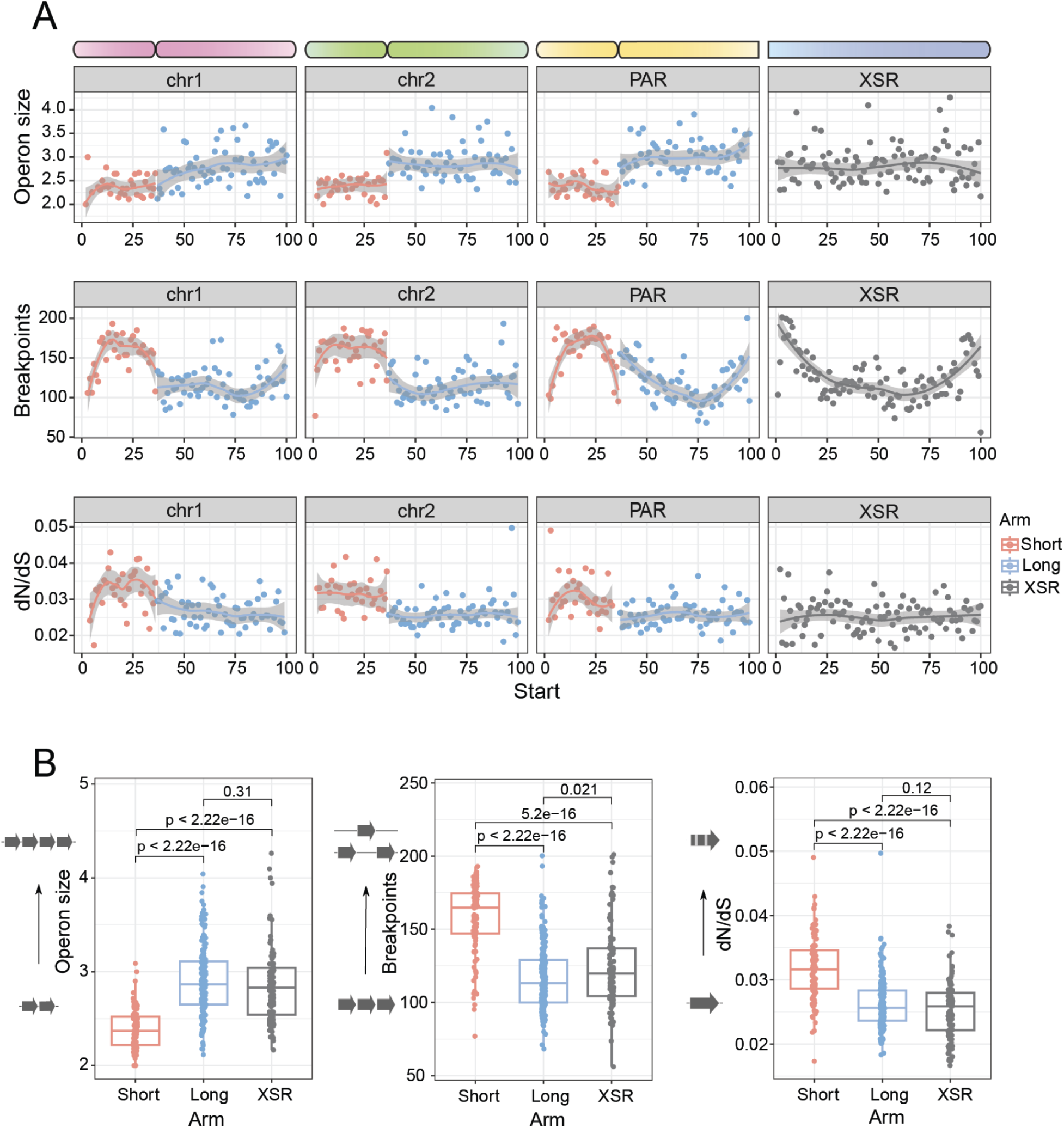
Genome-wide patterns of genomic feature density. A) The mean values for various genomic features (Y axis) versus chromosomal location by percentage of each chromosome’s length (X axis). Two regions of the chromosomes exhibit characteristic differences in feature distribution: the first difference can be seen between short and long chromosome arms, and the second difference is between the centres and edges of chromosome arms. B) Short and long arms exhibit significant differences in operon size, the number of breakpoint regions, and dN/dS ratios (Wilcoxon rank-sum test).

### Evolutionary framework of the unprecedented genome scrambling in *O. dioica* cryptic species

Considering the remarkable similarity of morphology and ecology of the *O. dioica* cryptic species studied here (Masunaga et al. 2022), the extent of scrambling that we have observed in the chromosomal assemblies appears to be unprecedented. In order to relate the rate of scrambling to evolutionary distance, we estimated a species tree and divergence times for *O. dioica* using orthologues common to chordates (Figure 6, Supplementary Table 2). The phylogenetic tree strongly supported the existence of three independent lineages of *O. dioica* cryptic species, which were estimated to have shared a last common ancestor ∼25 million years ago (Mya). This divergence represents the split between the subtropical North Pacific and other lineages, and a more recent divergence time of around ∼7.3 Mya was estimated for the split between the temperate North Pacific and North Atlantic lineages. Using these divergence time estimates, we calculated that the rate of breakpoint accumulation for *O. dioica* lies between 6 and 25 breakpoint regions per Mbp per million years (Figure 7A, D). By comparison, the estimated rate of breakpoint accumulation for *Ciona intestinalis* and *Ciona robusta* – a pair of tunicate ascidians with highly similar morphology – was ∼0.3 breakpoint regions per Mbp per million years, or an order of magnitude lower than *O. dioica* (Figure 7D). The difference in the degree of scrambling between animal species is immediately apparent by comparing the differences in collinearity between *Oikopleura* and *Ciona*; long segments of *Ciona* genomes maintain collinearity between species. A level of synteny dissolution comparable to *O. dioica* can only be observed between *Ciona* spp. in comparisons of highly morphologically divergent species with far older divergence times (∼100 Mya). We also compared the rates we estimated for *O. dioica* against *Drosophila* spp., as an example of a group of animals with similarly short generation times, high speciation rates, and which have been reported to exhibit a considerable degree of genome scrambling, which thus presumably have been affected by similar evolutionary dynamics (Suvorov et al. 2022). We used D. busckii for the split between the *Sophophora* and *Drosophila* subgenus (∼47 Mya), *D. subpulchrella* for the split between the *suzukii* and *melanogaster* subgroups (∼13 Mya), *D. yakuba* for a speciation within the *melanogaster* subgroup (∼7 Mya), and estimated between 0.5 and 3 breakpoint regions per megabase per million years. Strikingly, based on these metrics, all pairs of *Oikopleura* species showed a distinctly greater rate of scrambling than any other species (Figure 7D).

**Figure 6.**
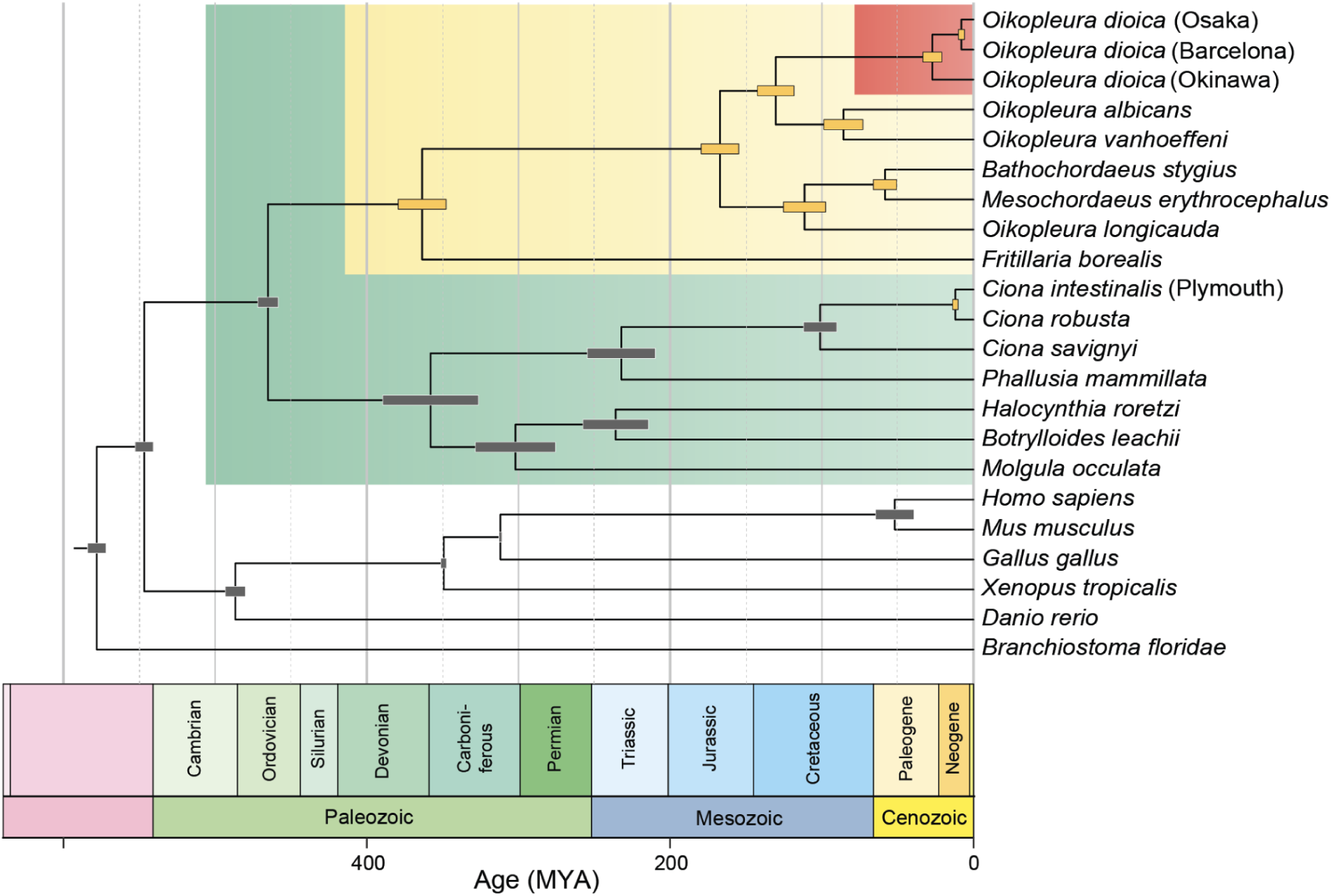
Time-scaled phylogenetic tree including several appendicularians, tunicates, and vertebrates including *O. dioica*. The different clades of *O. dioica* cryptic species were estimated to have shared a common ancestor ∼25 Mya (95% HPD: 12-41 Mya). The populations from the North Atlantic and North Pacific were estimated to have diverged more recently ∼7 Mya (95% HPD: 5-10 Mya).

**Figure 7.**
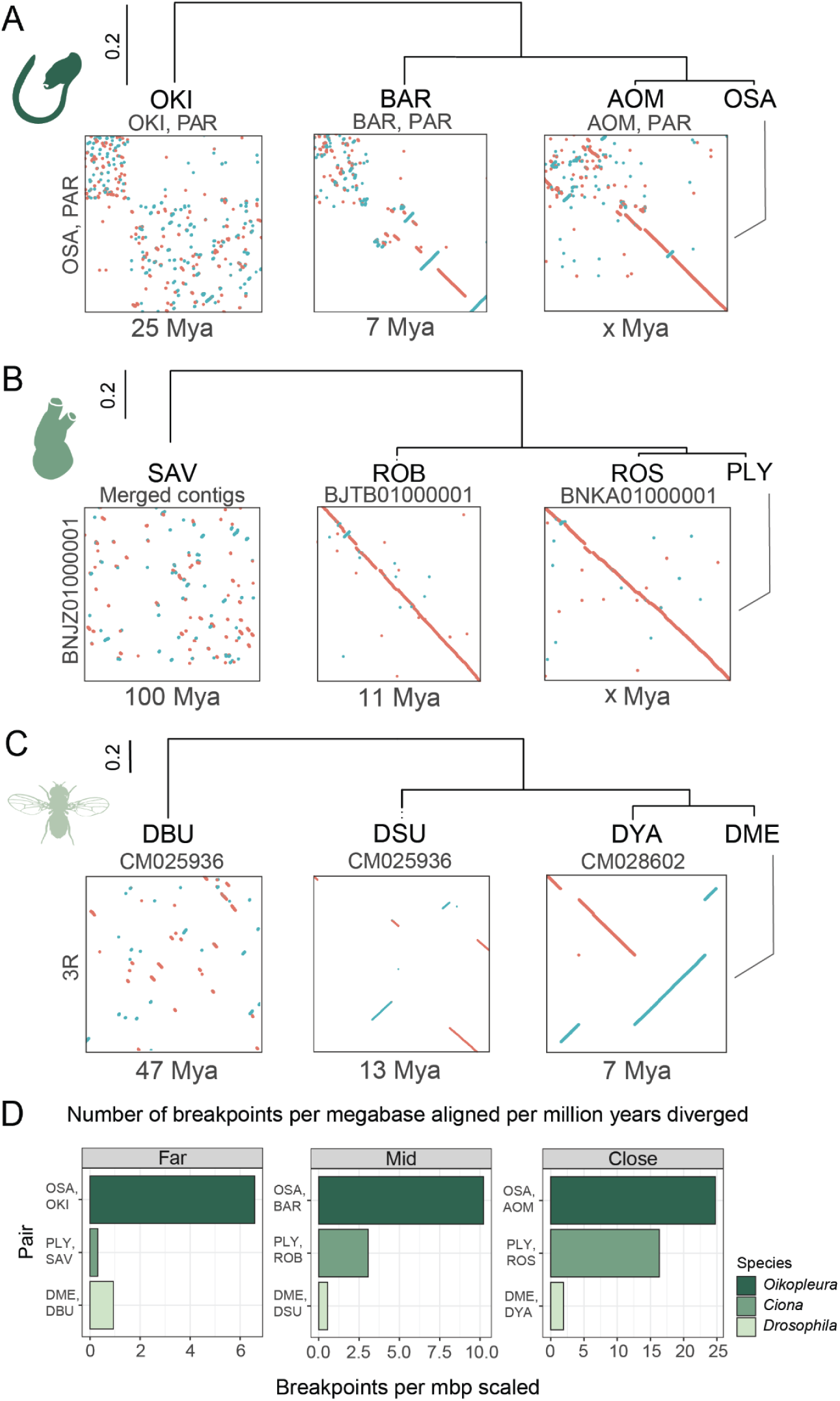
A) Genome scrambling in 10 × 10 Mbp regions for pairs of genomes in *Oikopleura* (A, dark green), *Ciona* (B, green), and *Drosophila* (C, light green). D) The number of breakpoint regions per megabase aligned per million years diverged for different animal clades. Abbreviations: OKI: Okinawa, BAR: Barcelona, OSA: Osaka, AMO: Aomori, SAV: *Ciona savignyi*, ROB: *C. robusta*, ROS: *C. intestinalis* (Roscoff), PLY: *C. int.* (Plymouth), BUS: *Drosophila buskii*, DSU: *D. subpulchrella*, DYA: *D. yakuba*, DME: *D. melanogaster*.

## DISCUSSION

We report extensive scrambling of gene order in three species of *Oikopleura dioica* that are so morphologically similar to each other that they are classified as a single species (Masunaga et al. 2022). Our study design utilising 6 genome assemblies ensures that our findings are supported by at least two inter-population comparisons, and each of these comparisons are consistent with the comparison to a second genome from the same species. This design, combined with the surprisingly recent divergence times estimated for *O. dioica*, allowed us to study genome scrambling on a finer time scale than previous studies (Drosophila 12 Genomes Consortium et al. 2007; Albertin et al. 2022; Hane et al. 2011). Further, the chromosomal resolution of our genome assemblies was essential, as it revealed consistent patterns of evolutionary rate heterogeneity within and between chromosomes in *Oikopleura*. We anticipate that ongoing efforts to produce high-quality, high-contiguity genome assemblies may prove useful to assess the prevalence of this phenomenon in other clades.

The sex chromosomes of *O. dioica* have 3 non-interacting Hi-C domains: short arms, long arms, and sex-specific regions. The X-specific region is comparable in size to long arms and displays similar features in terms of scrambling, alignment lengths, GC content, and gene/repeat density. The assemblies of the Okinawa and Barcelona genomes suggest that the sex-specific regions and the pseudoautosomal region long arm are separated by an rDNA locus. Thus, the limited interaction between the sex-specific and pseudoautosomal regions could be related to the division of the sex chromosome’s long arm by the rDNA locus, acting similarly to the centromeres that divide short and long arms elsewhere.

In the *Oikopleura* genomes we analysed, most genomic features exhibit bimodal distributions, with short and long modes (Figure 5). This global bimodality reflects a general tendency for short arms to scramble and evolve faster than long arms. We also observed consistent differences between the edges and centres of arms. As such, it is possible that the differences in genomic features between short and long arms could be explained by long arms possessing proportionally greater central regions. Further research will be needed to determine if there is a causal relationship, and the direction of the relationship, between domain length and the rate and extent of scrambling.

Genome alignments can associate overlapping query regions to multiple target regions, for instance because of paralogues or repeat elements. In their seminal work comparing the human and mouse genomes, Kent et al. reduced these many-to-many alignments networks into “chains” of ordered one-to-many alignments allowing for local inversions (Kent et al. 2003). While this approach may be robust for identifying large-scale syntenies between two genomes, it disregards true breakpoint regions that flank inversions. For this reason, we focused on the use of optimal one-to-one alignments (Frith and Kawaguchi 2015), and defined that a change in alignment strand breaks collinearity, but not synteny. Future developments may allow us to more efficiently utilise the many-to-many alignments – for instance, to provide robustness against whole-genome duplications, which did not happen in *Oikopleura dioica* – or to better resolve the causes of non-alignment, which could stem from transposon activity or segmental duplications and deletions. However, while our approach is primarily based on nucleotide alignments, the protein orthologue set computed from our genome annotations fully supports the fact that nucleotide sequence scrambling has resulted in scrambling of gene order. Thus, our nucleotide-based approach is an efficient way to study scrambling in the absence of gene annotations.

Our strict definition of collinearity increases the number of identified breakpoint regions in comparisons of closely-related populations compared to more distantly-related populations, because greater alignability of repetitive elements results in greater detection of transpositions (Figure 6). As well, the movements of repetitive elements that result in breakpoint regions in intraspecific comparisons may represent transient or allelic variations, which could be considered qualitatively different than breakpoint regions identified in inter-specific comparisons, which might represent ancient or fixed rearrangements from early in the speciation process. Whether or not the movements of repetitive elements have important functional consequences, they do represent break points requiring the breakage and repair of DNA, which means that the movement of repetitive elements necessarily represents at least one true molecular breakpoint. Since our methods rely upon alignments and therefore the limits of alignability, we have mostly focused our study on comparisons of distantly-related *O. dioica* lineages. Further investigations will be needed to better account for the movements of repetitive elements within and between the different *O. dioica* cryptic species.

Genes and their constituent exons are, perhaps unsurprisingly, strongly conserved between species of *O. dioica*. Within genes, introns typically occupy bridge regions within collinear alignments, and by definition, bridge regions do not have a corresponding segment in a pair of genomes. This observation is consistent with the high intron turnover reported by Edvardsen et al. (2004). Surprisingly, operons are rarely conserved between species. At first glance, it is puzzling that such extensive genome scrambling has occurred in an organism where a significant fraction of the transcriptome relies upon, at minimum, a segment of DNA containing a promoter and multiple collinear genes. However, two properties of *O. dioica* operons could be related to the observation that operons are not strictly maintained: first, the expression levels of operonic genes are not strongly correlated (Danks et al. 2015), and second, the functional categories of operonic genes are not necessarily correlated. Together, these observations suggest that the operon structure in *O. dioica* need not solely or primarily be related to the regulation of transcription. On the contrary, the presence of the operon transcriptional system could act to decrease the necessity for genes to retain their own promoters, by allowing them to freely insert into other operons with their own transcriptional machinery. Indeed, we identified several lines of evidence suggesting that operon-switching can occur. As such, an operon system such as it exists in *O. dioica* might in fact help to maintain gene expression in the context of genome scrambling.

At all levels of evolutionary distance, the widths of breakpoint regions are too large to easily identify the original molecular breakpoints of the DNA at the level of individual nucleotides. Two reasons may explain the lack of alignability in breakpoint regions: a) so many mutations accrue in these regions that they exceed the limits of detectable homology, or b) that repetitive elements were the target or the cause of the breaks. It is tempting to speculate that the loss of the canonical non-homologous end-joining (NHEJ) DNA repair pathway in *O. dioica* might have created synergies that act to promote scrambling. For instance, the alternative microhomology-based pathway (MMEJ), which was shown to be active in experimentally-induced lesions in *O. dioica* (Deng, Henriet, and Chourrout 2018), is slower than other repair mechanisms (Fu et al. 2021), which might allow for greater chromatin movement to occur before the repair of a double-stranded break. Cut-and-paste transposons that use the MMEJ pathway may also act as a source of microhomologies that could facilitate repair by MMEJ. The low repeat content of *O. dioica* genomes might therefore be a reflection of genomic instability that also causes scrambling. Although *O. dioica* genomes seem to be repeat-sparse, a relatively small number of interspersed repeats is sufficient to facilitate rearrangements through repair mechanisms such as homologous recombination. Scrambling in *O. dioica* also seems to correlate strongly with phylogenetic distance and divergence time. Parsimoniously, the mechanisms underlying scrambling are more likely to involve the gradual accumulation of rearrangements rather than the result of one or more dramatic lineage-specific rearrangement events. In any case, the mechanisms that contribute to scrambling in *O. dioica* do not seem to frequently involve significant gains or losses of gene content. Additional experiments are needed to elucidate the precise mechanisms underlying scrambling in *O. dioica*, perhaps through surveys of population-level genetic variation or monitoring the accrual of mutations in a controlled lab environment over time.

Accurately assessing divergence times is difficult, and molecular clock analyses are heavily influenced by their priors. In this study, we opted for relatively unconstrained priors, partly owing to a lack of suitable fossil taxa for calibrating the internal nodes of the appendicularian clade. A more restricted set of informative priors within this clade would allow more refined and precise divergence time estimates. The quality of the underlying phylogenomic data also has important consequences for divergence time estimation. Gene prediction in the appendicularia is difficult, particularly given the existence of operons and the use of non-canonical splice sites that complicate the generation of accurate gene models. Incorrect gene models could lead to orthologue misidentification and misalignment, which will inflate divergence time estimates. Differences in substitution rates between lineages (heterotachy), is also particularly problematic as *O. dioica* is among the fastest-evolving animals known (Berná, D’Onofrio, and Alvarez-Valin 2012). We attempted to minimise these effects by restricting our orthologue set to include only single-copy resolved orthologues, trimming gene alignments with HmmCleaner, removing low-information genes with MARE, and manually curating the orthologue set according to congruence with the highly-supported species tree (Simion, Delsuc, and Philippe 2020; Philippe et al. 2017). Given that many of these factors are likely to increase divergence estimates, as well as the relatively high estimate for the split between *Ciona robusta* and *Ciona intestinalis* (∼12 Mya; Bouchemousse, Bishop, and Viard 2016), we believe that our divergence time estimates for *O. dioica* are more likely to be overestimated than underestimated. Thus, although additional work is needed to confirm whether the level of scrambling exhibited by *O. dioica* is truly exceptional among animals, our results are consistent with an overall elevated rate of molecular evolution in *O. dioica*.

The species phylogeny we estimated suggests that at least 3 distinct species of *O. dioica* exist, which was corroborated by our analyses of marker genes (Masunaga et al. 2022) (Masunaga et al. 2022). Our molecular clock analysis suggests that the North Pacific and North Atlantic species shared a common ancestor ∼7 Mya, which diverged from the Okinawa group more than 20 Mya. These results are not yet reflected taxonomically. Ultimately, classifying these populations taxonomically will involve other facts and the needs of the community beyond this manuscript, taking into account factors such as the stability of names in the literature and practicality for field studies. Regardless, the genetic and environmental factors that might have contributed to cryptic speciation in this clade are unknown. It is tempting to speculate that the extreme rate of rearrangement in *O. dioica* could accelerate sympatric speciation through the formation of reproductively incompatible subpopulations within an area, even in marine environments lacking physical geographic boundaries. Further surveys of intrapopulation genetic variation are needed to validate this hypothesis.

Our discovery of the exceptional scrambling between species of *O. dioica* is corroborated by the fact that it has little to no detectable synteny remaining with the ascidian tunicate *Ciona intestinalis* (Denoeud et al. 2010), nor with amphioxus, which has previously been used as a proxy for the ancestral linkage groups of all chordates (Simakov et al. 2022). By contrast, conservation of synteny dating back 400 million years has been identified in insects (Shah et al. 2012; Meisel, Delclos, and Wexler 2019). As no chromosome-scale assembly is yet available for other appendicularians, the speed of synteny loss in *O. dioica* compared to its relatives is unknown. Nevertheless, the position of *O. dioica* as one of the fastest-evolving metazoans – and indeed, perhaps one of the fastest-evolving eukaryotes – makes it a fascinating case study in the evolution of synteny.

## CONCLUSIONS

We report that the genome of the planktonic tunicate *Oikopleura dioica* has rearranged thousands of times in ∼25 million years without detectable differences in morphology or ecology, making it the fastest-evolving animal genome yet described. The pace of rearrangement differs within and between chromosome arms, with a faster pace observed in short arms. Further work will be required to investigate the roles that the loss of the canonical NHEJ DNA repair pathway and the existence of gene operons may have had in promoting genome scrambling. The extent of scrambling that happened naturally in *Oikopleura dioica* may also provide cues on how to design synthetic genomes with flexible gene order. Finally, we expect that the pool of natural diversity is not yet exhausted, as there are no genomic studies from *Oikopleura* sampled from the Southern Hemisphere.

## MATERIALS AND METHODS

### Public data

We downloaded the OKI2018_I69 assembly (Bliznina et al. 2021) from doi:<u>10.1186/s12864-021-07512-6</u>, and removed the unplaced scaffolds. We downloaded the OdB3 (Bergen, Norway) assembly (Denoeud et al. 2010) from https://www.genoscope.cns.fr/externe/Download/Projets/Projet_HG/data/assembly/unmasked/O <u>dioica_reference_v3.0.fa</u>.

### Sampling, genome sequencing, genome assembly, and scaffolding

We extracted high-molecular weight DNA from one individual (“Bar2”) from the Barcelona laboratory strain (Martí-Solans et al. 2015) using a modified salting-out protocol (Masunaga et al. 2022), sequenced it on MinION sequencer Mk1B (Oxford Nanopore Technologies) using a SQK-LSK109 kit (ONT) following the manufacturer’s instructions and basecalled it with the Guppy software (ONT) version 4.4.2 using the Rerio model res_dna_r941_min_crf_v031 (https://github.com/nanoporetech/rerio). The shortest reads were discarded until remaining data reached 7 × 10^9^ nt, using the filtlong software (https://github.com/rrwick/Filtlong), resulting in a read N50 higher than 30,000 nt. We then assembled the genome using the Flye software (Kolmogorov et al. 2019) version 2.8.2-b1689 with the --min-overlap 10000 parameter using a custom Nextflow pipeline (doi:10.5281/zenodo.6601989). To ensure one-to-one correspondence between assemblies, we removed alternative haplotype sequences using the purge_dups tool (Guan et al. 2020). However, as a single removal step was not efficient enough, we employed an iterative approach where haplotigs were flagged with purge_dups, reads mapped uniquely to contigs with LAST and last-split, and then reads mapping to purged haplotigs were removed before restarting the whole assembly process. Iterations were stopped after purge_dups stopped discovering haplotigs, and an assembly was selected that provided the best tradeoff between contiguity (which typically increased during the first iterations) and a low number of duplicated single-copy orthologues. The contigs were then polished with Pilon 1.22 (Walker et al. 2014) using short-read sequences from the same individual (<u>PRJEB55052</u>), and scaffolded using Hi-C data from the Bergen line at tailbud stage (SRR14470734) using Juicer (Durand et al. 2016) and 3D-DNA (Dudchenko et al. 2017), like in Bliznina et al. 2021.

We sequenced the Kume and Aomori genomes using single animals isolated from wild populations (Masunaga et al. 2020) with the same method except that we basecalled with Guppy version 5.0.11 and the Guppy model dna_r9.4.1_450bps_sup, and used Flye version 2.8.3-b1763 with parameters --min-overlap 10000 --extra-params assemble_ovlp_divergence=0.04,repeat_graph_ovlp_divergence=0.04,read_align_ovlp_diverge nce=0.04,max_bubble_length=800000,use_minimizers=1,minimizer_window=5, and that no scaffolding no polishing were performed.

We re-scaffolded the OSKA2016 genome (K. Wang et al. 2020) by merging scaffolds that were overlapped by long contigs from single individual genome draft assemblies from the same laboratory strain (SAMEA6864573 and PRJEB55052). As a last resort, we arbitrarily merged some contigs to a chromosome arm based on synteny information. The resulting OSKA2016v1.9 assembly is described in more detail in https://github.com/oist/LuscombeU_OSKA2016_rescaffolding.

We sequenced genomes exclusively from male animals because they simplify the assembly of the sex-specific regions, which are single-copy in males.

For all genomes, we counted metazoan near-universal single-copy genes using the Benchmarking Universal Single-Copy Orthologs (BUSCO (Manni et al. 2021) tool version 5.2.1 and an AUGUSTUS model trained for annotating the OKI2018_I69_1.0 assembly (Bliznina et al. 2021; Hoff and Stanke 2019). While this version of BUSCO appears to have a lower detection baseline compared to the v3 series that we used for the OKI2018_I69 genome assembly (64% vs. 73%) (Bliznina et al. 2021), the completeness of our new assemblies is consistent with the score of the OKI2018_I69 genome assembly for which we previously demonstrated high completeness (Bliznina et al. 2021). Finally, we removed unplaced scaffolds from all chromosomal assemblies.

### Pairwise genome alignment and comparison

We aligned pairs of genomes using the same approach as in (Bliznina et al. 2021). In brief, we used the LAST software (Kiełbasa et al. 2011) to align a “query” genome to a “target” genome indexed with the YASS seed (Noé and Kucherov 2005) for long and weak similarities with parameters and a scoring matrix determined by the LAST-TRAIN software (Hamada et al. 2017) and filtered the resulting many-to-many set of alignment pairs with the last-split tool (Frith and Kawaguchi 2015), which searches for an optimal set of one-to-one local alignments, and finally removed alignments that include a significant amount of masked sequences with the last-postmask tool (Frith 2011). We parallelised this process in a Nextflow (Di Tommaso et al. 2017) workflow available at (https://github.com/oist/plessy_pairwiseGenomeComparison/tree/v5.1.0). To load the alignment coordinates in the R environment for statistical computing (R Core Team 2023), we wrote a package called “GenomicBreaks” (https://oist.github.io/GenomicBreaks/) using core Bioconductor libraries (Lawrence et al. 2013). In this package the strand randomisation index is computed for each chromosome by subtracting the total length of opposite-strand alignments to the total length of same-strand alignments and dividing the result by the total length of the aligned regions so that a value of 1 indicates that all alignments are same-strand, and a value of zero indicates that overall the orientations appear to be random. The average of the values obtained on each chromosome is then computed and weighted by the length of the chromosomes. For the computation of the breakpoint and bridge regions, we used the strictest definition of collinearity, where it is interrupted by inversions (changes of alignment strand) and translocations (presence of one extra aligned region in one genome only) of any length. A copy of the software and the alignment files is archived on Zenodo (doi:10.5281/zenodo.6601989). A rendering of the R vignettes that we used to produce these visualisations are available at https://oist.github.io/LuscombeU_OikScrambling/ and compiled as an interactive notebook of vignettes in Supplementary data 1.

Pairwise comparison between the *O. dioica* genomes produced in this work and the *O. vanhoeffeni*, *O. longicauda,* and *O. albicans* genomes (Naville et al. 2019) were loaded in the CNEr package (Tan, Polychronopoulos, and Lenhard 2019) to define conserved non-coding elements with a window size of 50 and an identity threshold of 48 (doi:10.5281/zenodo.6601989).

### Repeat masking and gene annotation

For each genome, a custom library of the repetitive elements was created by merging outputs of three different software: RepeatModeler (Flynn et al. 2020) version 2.0.1, MITE-Hunter (Han and Wessler 2010) version 11–2011, and SINE_Finder (Wenke et al. 2011), which were used as input for RepeatMasker (Smit, Hubley, and Green 2013) version 4.1.0. The repeats identified by homology searches were soft-masked in each assembly.

Gene models were predicted using AUGUSTUS (Stanke et al. 2006) v3.3.3 using the species model trained for the OKI2018_I69 *O. dioica* (Bliznina et al. 2021). In order to produce more accurate annotations, transcripts aligned to genomes with BLAT (Kent 2002) version 36 were used as “hints”. In cases where an assembled transcriptome was not available, data from related individuals was used. In particular, the transcriptome assembly generated by Wang et al. (K. Wang et al. 2015) was used for predicting genes in both OSKA2016v1.9 and AOM-5-5f genomes, while the transcriptome assembly generated by Bliznina et al. was used for re-annotation of OKI2018_I69 genome and annotation of the KUM-M3-7f genome. A Barcelona transcriptome assembly was used for gene prediction in the Bar2_p4 genome. The parameter “--allow_hinted_splicesites” was used with AUGUSTUS to allow the prediction of non-canonical splice sites (GAAG, GCAG, GGAG, GTCG, GTAA). Operons were annotated for each species, defining an operon as a set of collinear genes separated by an intergenic distance of at most 500 base pairs.

### Orthologue identification

Gene orthology was reconstructed using OrthoFinder (Emms and Kelly 2015; 2019) version 2.5.4 based on 26 proteomes spanning three subphylums of chordates (data doi:10.5281/zenodo.6601989). To improve orthology assignment within *O. dioica*, multiple tunicate species were included as recommended in the OrthoFinder tutorials (https://davidemms.github.io/). Gene predictions for six appendicularian genomes from Naville et al. (2019) and two geographically-distinct *Ciona intestinalis* genomes (Plymouth and Roscoff; Satou et al. 2021) were computed using a similar approach to *O. dioica*, including repeat-masking followed by gene prediction with AUGUSTUS version 3.3.3. Gene prediction used either the *O. dioica* or *Ciona* model, as other species lack publicly available gene annotations. The proteomes of other species were downloaded from Uniprot: *Branchiostoma floridae* (UP000001554), *C. intestinalis* type “A” (*robusta*, UP000008144), *C. savignyi* (UP000007875), *Danio rerio* (UP000000437), *Xenopus tropicalis* (UP000008143), *Gallus gallus* (UP000000539), *Mus musculus* (UP000000589), *Homo sapiens* (UP000005640). Four more tunicate species were included from the Aniseed database: *Botrylloides leachii*, *Halocynthia roretzi*, *Molgula oculata*, *Phallusia mammillata*. To remove redundancy in the dataset, protein sequences were clustered at 100% identity using CD-HIT (Li and Godzik 2006) version 4.8.1. Alternative haplotypes were removed from the Bergen *O. dioica* proteome. and only the longest isoforms per gene were used for the analysis. OrthoFinder was run with the parameters -M msa -T raxml-ng with the following fixed species tree to ensure that *O. dioica* sequences fall within the oikopleurid branch:

(((Danio_rerio,(Xenopus_tropicalis,((Mus_musculus,Homo_sapiens),Gallus_gallus))),(((Molgula_oculata,(Halocynthia_roretzi,Botrylloides_leachii)),((Ciona_savignyi,((C_intesinalis_P,C_intestinalis_R),Ciona_robusta)),Phallusia_mammillata)),(Fritillaria_borealis,((Oikopleura_longic auda,(Mesochordaeus_erythrocephalus,Bathochordaeus_sp)),((Oikopleura_vanhoeffeni,Oikopl eura_albicans),((KUM-M3-7f,OKI2018_I69),((Bar2_p4,OdB3),(AOM-5-5f,OSKA2016v1.9)))))))), Branchiostoma_floridae);

The Hox protein sequences of the Bergen population were used as reference Hox sequences for *O. dioica* (H.-C. Seo et al. 2004). In general, Hox genes were assigned appropriate orthogroups by OrthoFinder, although Hox11 could not be identified within the Barcelona proteome and the Hox9 model for Osaka had not been spliced appropriately. Regardless, the identities of these genes were confirmed by alignment with the Bergen sequence as well as multiple sequence alignment with all orthologous family members followed by tree estimation with IQ-TREE (Nguyen et al. 2015) version 1.6.12.

### Phylogenomics and divergence time estimation

A set of single-copy orthologue sequences was extracted from the results of the OrthoFinder run, selecting proteins that were shared by 10 or more of the 26 species, yielding 555 orthologue candidates. Each orthologue set was aligned using PRANK (Löytynoja 2014) v.170427 then trimmed with HmmCleaner (Di Franco et al. 2019) and a gene tree was estimated with RAxML (Stamatakis 2014) version 8.2.4, with 100 rapid bootstraps and a gamma model of rate heterogeneity with automatic model selection using PROTGAMMAAUTO. Each gene tree was compared to the later species tree with the ete3 toolkit (Huerta-Cepas, Serra, and Bork 2016). A supermatrix was constructed and gene information content was assessed with MARE (Meyer, Meusemann, and Misof 2011) v0.1.2-rc, which reduced the number of genes to 177. The alignment supermatrix generated from these 177 genes was used to estimate a species tree with RAxML using 100 rapid bootstrap replicates, the gamma model of rate heterogeneity, and automatic model selection for each gene as separate partitions. To estimate divergence times, BEAST1 (Suchard et al. 2018) v1.10.4 was used with the beagle library (Ayres et al. 2012) with the following parameters: the birth-death tree density model (Gernhard 2008), a linked random local clock model (Drummond and Suchard 2010), an unlinked gamma-distributed rate heterogeneity with 4 categories for each partitioned gene, and the CTMC scale reference prior model (Ferreira and Suchard 2008). In order to estimate only divergence times, the tree topology was fixed to the species tree estimated by RAxML. Where possible, the divergence time estimates published by (Delsuc et al. 2018) using their LN CAT-GTR + Γ_4_ model were used as normally-distributed priors on our tree with matching mean and standard deviation. Each node that did not correspond between the two studies, including the appendicularian proteomes that we annotated, uses uniformly-distributed priors with a maximum age as the age of the tunicates. The only exception was a normally-distributed prior for the split between *Ciona intestinalis* and *Ciona robusta*, which used the value reported by Bouchemousse et al. (2016). To ensure the models had converged, Tracer (Rambaut et al. 2018) was used v1.7.2; Rambaut et al. 2018, PMID 29718447), and further, three replicate analyses were performed using these parameters, taking the last 100M steps post-convergence for calculating statistics. The final re-sampled, combined metrics are reported in (Supplementary). The maximum clade credibility tree with node heights summarised to the median is depicted in Figure 5, using the replicate with the best marginal likelihood estimated by generalised stepping-stone sampling. The R libraries ggtree (Yu 2020) version 3.2.1, treeio (L.-G. Wang et al. 2020) version 1.18.1, and deeptime (Hoffmann et al. 2022) version 0.2.2 were used for tree visualisation.

### dN/dS estimation

To generate dN/dS estimates for *O. dioica* genes, single-copy orthologous proteins common to all 6 *Oikopleura dioica* proteomes were assessed. Each orthologous protein was aligned using PRANK, then unreliable sites were trimmed using GUIDANCE2 algorithm (Sela et al. 2015) v2.02. Protein alignments were converted to codon alignments using PAL2NAL (Suyama, Torrents, and Bork 2006) v14.1. Trees were estimated for each orthologue using RAxML with the PROTCATAUTO model and 100 rapid bootstraps. Then, dN/dS values were estimated using the CODEML program of the PAML package (Z. Yang 1997; Ziheng Yang 2007) version 4.9j using several different parameter combinations. Different parameters had little overall effect on the values or the interpretation of the dN/dS measurements, providing some validation for our method (data not shown). Ultimately, the dN/dS values estimated from a global comparison of all species (i.e., a single dN/dS estimate per orthologue) were used in Figure 4 and elsewhere.

## AUTHOR CONTRIBUTIONS

● Conceptualization: CP, NML
● Data curation: AB, CP, MJM, PN
● Formal Analysis: AB, CP, EMT, MJM, PN
● Funding acquisition: CC, EMT, NML
● Investigation: AB, AM, AWL, CC, CP, CW, GSS, JG, MFT, MJM, MSRP, NML, PN, TO, YT
● Methodology: AFR, CP, CW, MJM, PN, VR
● Project administration: CP, NML
● Resources: HN, EMT, CC
● Software: CP, MJM
● Supervision: CP, EMT, NML
● Validation: CP, MJM
● Visualization: AB, AM, CP, MJM
● Writing – original draft: AB, CP, MJM, NML
● Writing – review & editing: CP, EMT, MJM, PN, CC, NML

## ACKNOWLEDGEMENTS

We would like to thank the DNA Sequencing Section and the Scientific Computing and Data Analysis Section of the Research Support Division at OIST for their support, Ferdinand Marlétaz and Tom Bourguignon for critical comments, Cristina Frías Lopez for for initial bioinformatic support on the Barcelona genome assembly, and Atsuo Nishino for providing the Aomori samples. This work was supported by OIST core funding and in part by grant NFR-FRIBIO 204891/F20 to E.M.T. from the Norwegian Research Council. M.J.M. gratefully acknowledges funding from the Japan Society for the Promotion of Science as a JSPS International Research Fellow (Luscombe Unit, Okinawa Institute of Science and Technology Graduate University). CC was funded by BFU2016-80601-P and PID2019-110562GB-I00 from the Spanish Ministerio de Ciencia e Innovación and by 2021 SGR00372 AGAUR, Generalitat de Catalunya; VR by 2017BP00139 AGAUR, Generalitat de Catalunya and 2019IRBio001 from IRBio, Universitat de Barcelona; GSS by FPU18/02414 fellowship from Ministerio de Educación y cultura M.P.-C. by colaboración-2015/16, MFT by a PREDOC2020/58 fellowship from Universitat de Barcelona; AFR by MS12 Margarita Salas from Ministerio de Universidades (Spain). The sequencing of the *O. dioica* genome from Barcelona has been done under the Catalan Initiative for the Earth Biogenome Project.

