## Supplementary Figures for "Extreme genome scrambling in cryptic *Oikopleura dioica* species"

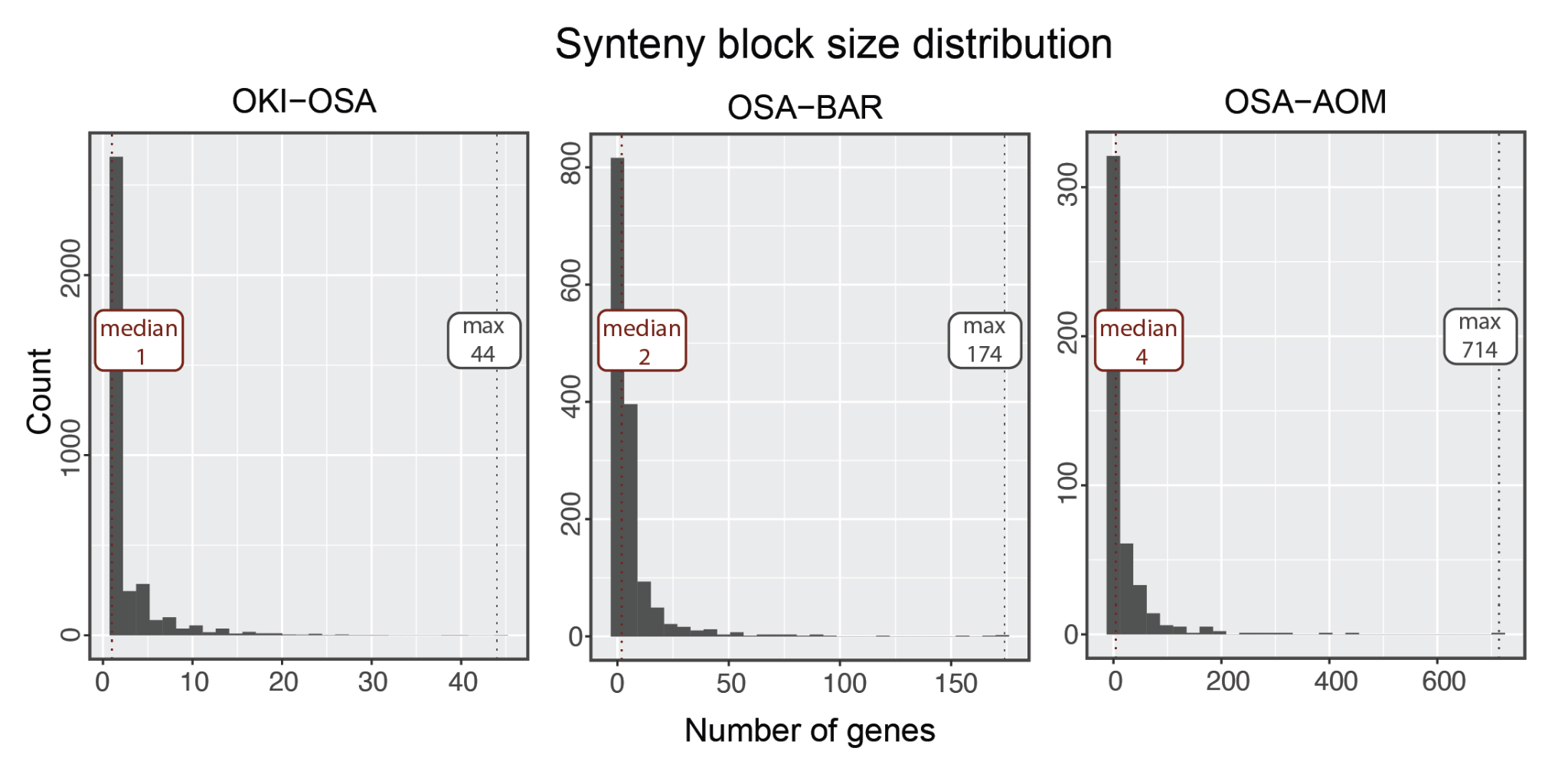

**Supplemental Figure S1:** Histogram of the number of orthologous genes per syntenic region in pairs of genomes.

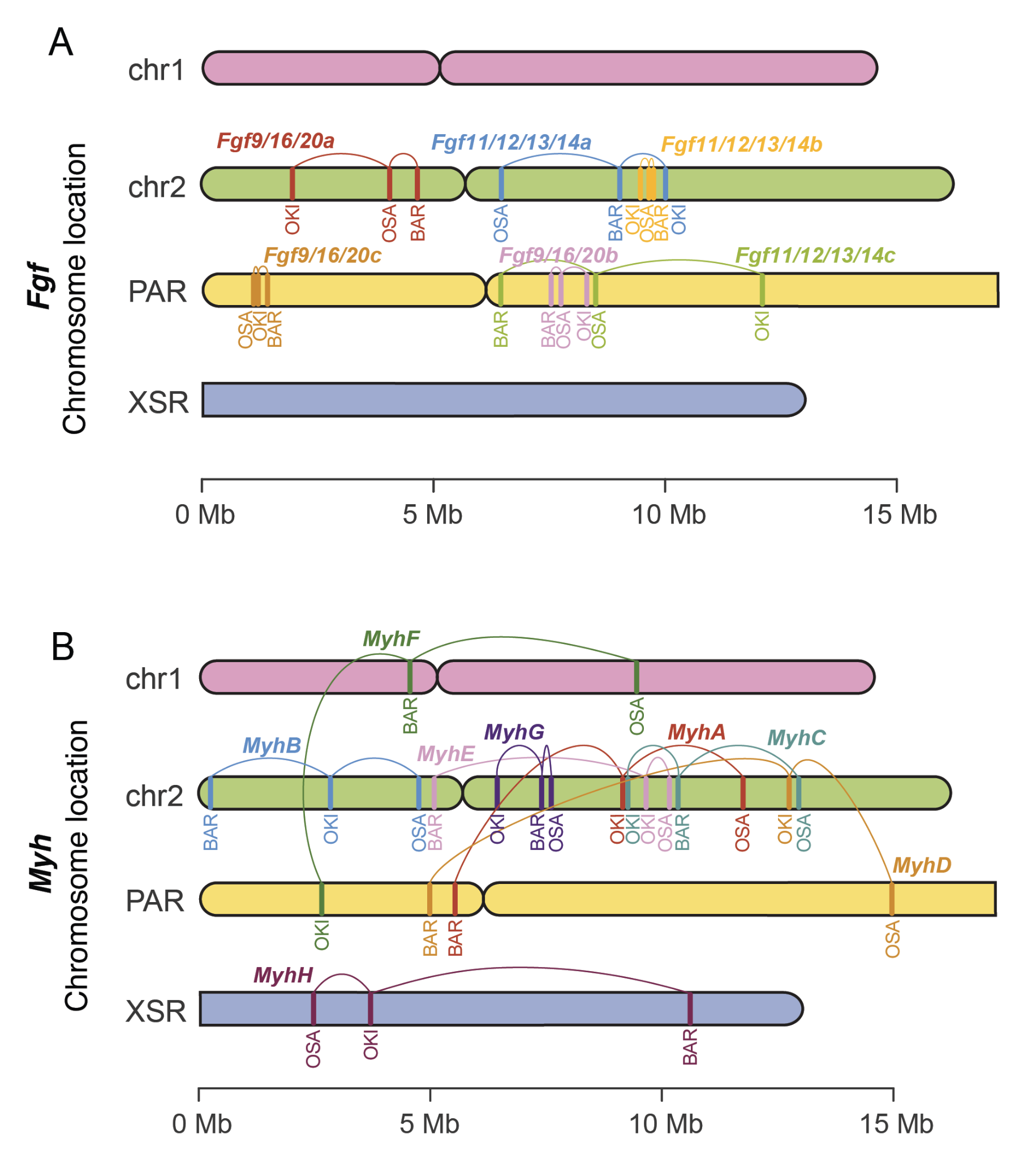

**Supplemental Figure S2:** Comparative chromosome mapping of the Fgf (A) and Myh genes in the genomes of *O. dioica* from Osaka (OSA), Barcelona (BAR) and Okinawa (OKI).

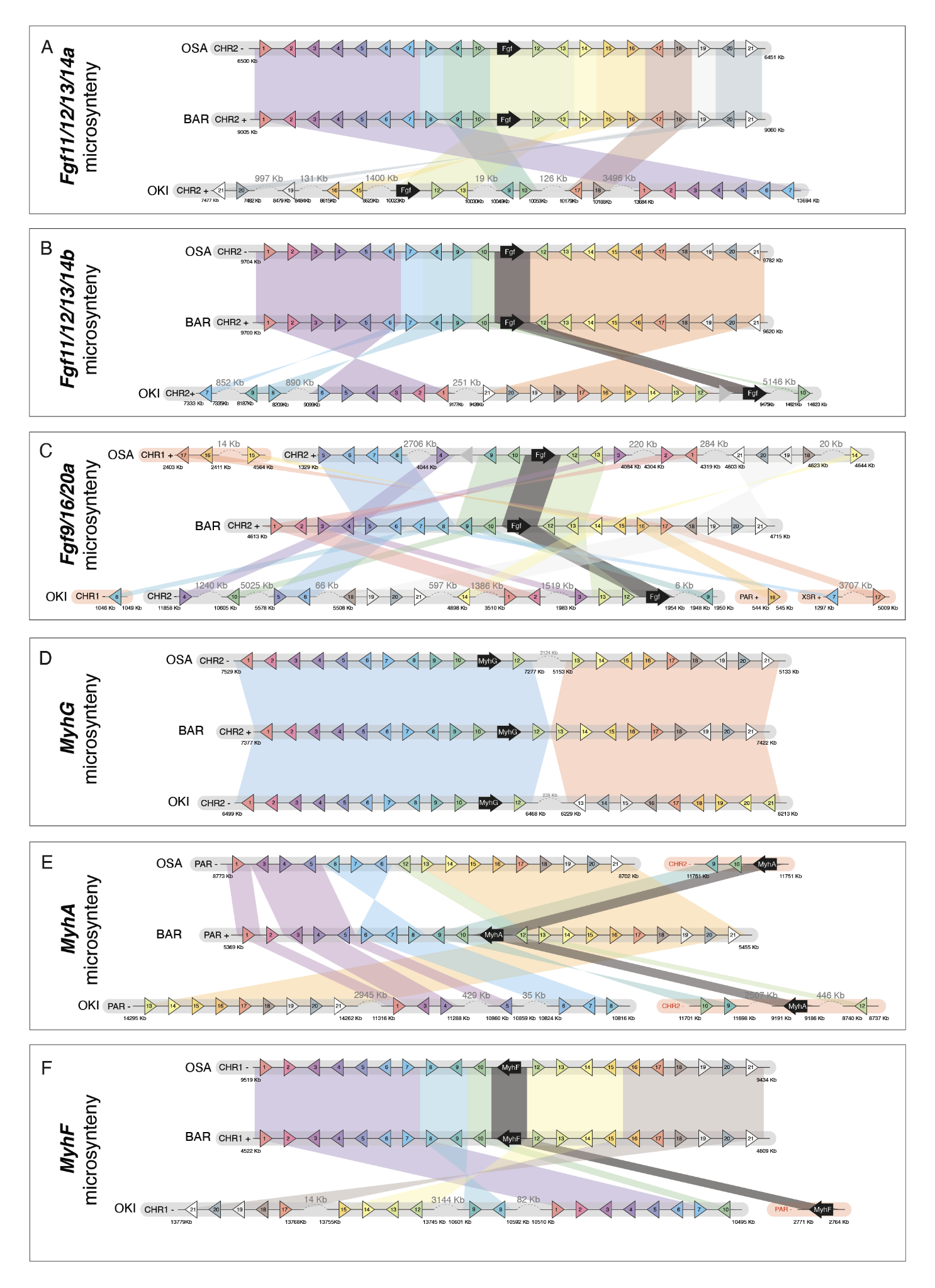

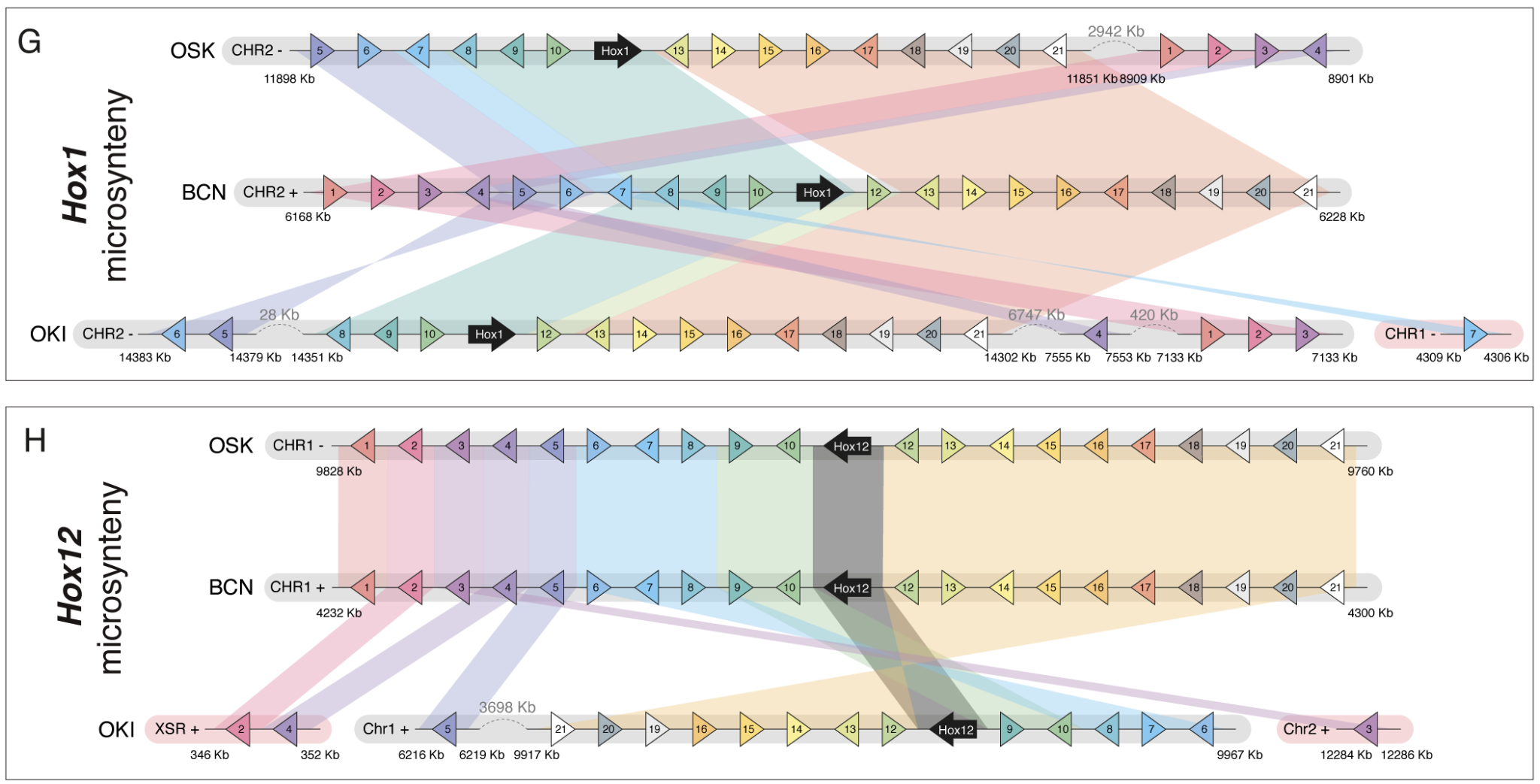

**Supplemental Figure S3:** Comparative microsynteny analysis of loci surrounding *Fgf* and *Myosin* gene family members in *O. dioica*. A: *Fgf11/12/13/14a*; B: *Fgf11/12/13/14b*; C: *Fgf9/16/20*) and *Myh* (D: *MyhG*; E: *MyhA*; F: *MyhF*). Species names are shortened as follows: Osaka (OSA), Barcelona (BAR) and Okinawa (OKI).

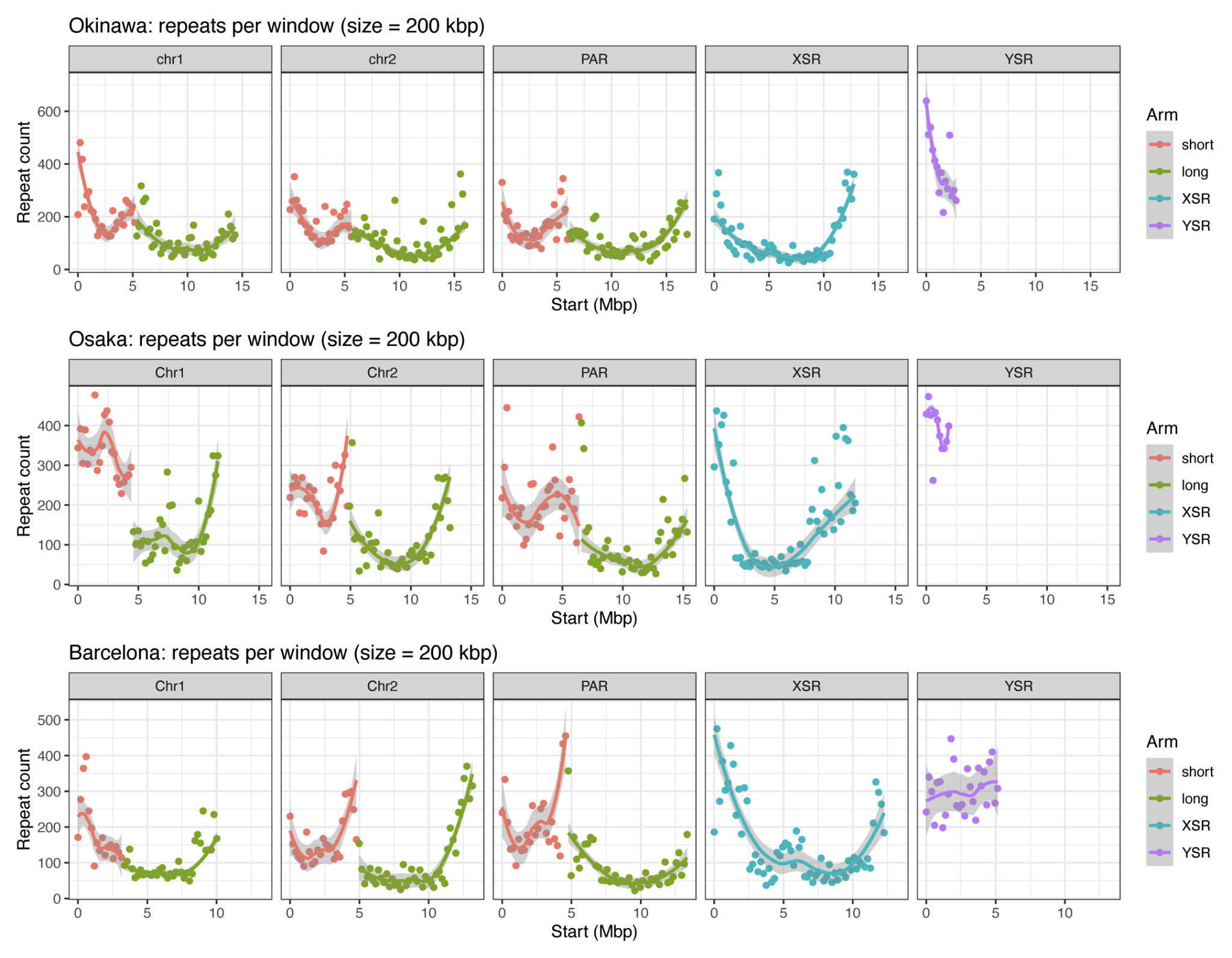

**Supplementary Figure S4**: Repeat density in *Oikopleura* genomes. The result for the Okinawan genomes was originally reported in (Bliznina et al. 2021), but is plotted here to facilitate comparisons with *O. dioica* from Osaka and Barcelona.
